# Searchlight Optimization Using Representational Similarity Analysis for Subject-Level Voxel Selection in Emotional State Decoding

**DOI:** 10.64898/2026.06.16.729835

**Authors:** Xuelei Wang, Jana Zweerings, Michael Lührs, Fengyu Cong, Klaus Mathiak, David E.J. Linden, Rainer Goebel, Assunta Ciarlo, David M. A. Mehler

## Abstract

Identifying informative voxels is a critical, yet challenging step in functional magnetic resonance imaging (fMRI), particularly for multivariate analyses involving multiple related conditions. Existing approaches often rely on predefined regions of interest (ROIs) or activation-based criteria, which may be insufficient for capturing fine-grained representational differences. This challenge becomes particularly relevant in experimental settings and interventions such as neurofeedback training, where voxels are not only measured as neural responses but also used as targets for intervention based on their previously observed activity patterns. In this study, we propose a subject-level searchlight optimization framework that integrates voxel-wise general linear model (GLM)–based univariate analysis with representational similarity analysis (RSA)–based multivariate refinement to identify voxels that are both task-relevant and condition-sensitive. To enhance practical applicability, the framework further incorporates a data-driven hyperparameter tuning step based on Bayesian optimization, enabling efficient identification of high-performing configurations from small pilot datasets, with consistent performance when applied to larger samples. The proposed framework was evaluated using an emotion imagery fMRI dataset with four affective conditions. Results demonstrate that the multivariate refinement improves alignment between empirical and target representational structures compared with univariate selection alone. Compared with a classifier-based voxel selection approach, the RSA-based approach better preserves the representational geometry of emotional states while maintaining discriminative capacity. These findings highlight the effectiveness, efficiency, and robustness of the proposed RSA framework, providing a practical solution for identifying condition-sensitive voxels and supporting more precise multivariate investigation of affective brain states in multi-condition fMRI studies.

## 1. INTRODUCTION

Functional magnetic resonance imaging (fMRI) has been proposed as a powerful technique for examining how cognitive and affective processes are represented in the human brain (Alotaibi et al., 2025; Poldrack & Farah, 2015). However, these processes recruit only a subset of brain regions or even sparse clusters of voxels (Haxby, 2012; Haxby et al., 2001). The way these target regions are defined can strongly influence anatomical interpretation (Brett et al., 2002; Eickhoff et al., 2018; Glasser et al., 2016), statistical reliability (Tarhan & Konkle, 2020; Walther et al., 2016), and model generalization performance (Norman et al., 2006; Pereira et al., 2009). Therefore, selecting informative voxels or regions of interest (ROIs) has become a fundamental and challenging step in fMRI studies and in particular in multivariate paradigms (Haynes, 2015; Hebart & Baker, 2018), as it plays a critical role in optimizing analysis pipelines and improving method sensitivity for both fundamental and clinical research applications.

Traditional voxel selection approaches often rely on anatomically predefined ROIs or univariate activation contrasts. ROI-based approaches assume that voxels within the same anatomical region and across individuals share similar functional properties, which may overlook the fine-grained heterogeneity of cortical representations (Brett et al., 2002; Eickhoff et al., 2015, 2018; Frost & Goebel, 2012). Although univariate activation-based approaches relax anatomical constraints by selecting individual voxels based on amplitude differences, they remain insensitive to distributed multivariate patterns that carry substantial task-relevant information, particularly in multi-condition paradigms (Haxby et al., 2001; Kriegeskorte et al., 2006; Kriegeskorte & Bandettini, 2007; Norman et al., 2006). These limitations motivate the need for voxel selection approaches that can identify subject-specific and condition-sensitive representational geometry.

Multivariate pattern analysis (MVPA) provides a powerful framework for addressing these challenges by focusing on distributed activation patterns rather than treating voxels independently. Decoding-based approaches use classifier or regression models to identify multivoxel patterns that discriminate experimental conditions, offering greater sensitivity to subtle pattern differences (Haynes & Rees, 2006; Hebart & Baker, 2018; Norman et al., 2006). Similarity-based approaches characterize the geometric structure of multivoxel response patterns and enable direct comparisons with cognitive or computational models (Kriegeskorte et al., 2008; Nili et al., 2014; Shinkareva et al., 2013). Searchlight analyses can further extend these approaches by evaluating the corresponding multivariate metrics within spatially contiguous neighborhoods, allowing fine-grained mapping of distributed representations across the brain (Etzel et al., 2013; Kriegeskorte et al., 2006). The MVPA framework has been widely applied in fMRI studies (Bressler et al., 2024; Haynes, 2015; Weaverdyck et al., 2020), providing a solid methodological foundation for developing voxel selection approaches based on multivariate representational patterns.

Within this broad MVPA framework, similarity-based approaches—particularly those based on representational similarity analysis (RSA)—offer a strong conceptual match to theories in cognitive neuroscience. These theories characterize mental representations as continuous structures distributed across multidimensional psychological spaces rather than as discrete categorical states (Roads & Love, 2024). RSA naturally aligns with this perspective by quantifying the similarity across conditions and capturing continuous representational information (Kriegeskorte et al., 2008; Kriegeskorte & Kievit, 2013). This conceptual alignment has led to a broader interest in the use of RSA for studying representational organization across a range of domains, including motor encoding (Ejaz et al., 2015; Kolasinski et al., 2020; Yokoi et al., 2018), visual perception (Nordt et al., 2019, 2023), neural development (Nordt et al., 2021, 2023), social cognition (Popal et al., 2019), episodic memory (Chadwick et al., 2010), semantic cognition (Meersmans et al., 2020; Ren et al., 2025), and affective states (Kiyokawa & Hayashi, 2024). Furthermore, other studies have focused on meta functions such as cognitive control (Freund et al., 2021) and clinical translations to neurological (Fang et al., 2018; Fischer-Baum et al., 2017) and psychiatric populations (Kurihara et al., 2025; Matsumoto et al., 2023).

More recently, RSA has been explored for its potential in perturbing mental representations and neural representations: identified neural representations remain correlations and their causal role is unclear (Krakauer & Ramsey, 2026) without a precisely controlled intervention that allows to directly manipulate at this level (Weichwald & Peters, 2021). Real-time semantic fMRI neurofeedback has been suggested to address this challenge by perturbing neural representations via engaging mental representations (Goebel et al., 2024). First studies introduced a practical approach to identify informative voxels that encode mental representations by combining univariate activation filtering with RSA-based evaluation to achieve near–real-time voxel identification and demonstrated its feasibility for guiding neurofeedback (Ciarlo et al., 2022; Russo et al., 2021; Wang et al., 2026). However, these voxel selection approaches were implemented as fixed components of the experimental pipeline without systematic assessment of their performance or optimization of key hyperparameters such as searchlight configuration, selection thresholds, or mask size. In addition, their performance has not been clearly evaluated against widely used classifier-based voxel selection methods. Consequently, RSA-based voxel selection, despite its conceptual appeal, requires further investigation and formalization into an efficient, standardized optimization framework for multivariate fMRI analyses.

The present study introduces, to our knowledge, the first systematically fine-tuned RSA-based searchlight optimization framework for subject-specific voxel selection in multi-condition fMRI studies. Building on prior work that combines univariate filtering with RSA, our framework applies univariate preselection to reduce the search space and then evaluates local multivoxel patterns using RSA-based searchlights to identify voxels whose representational geometry aligns with a target model. To ensure practical applicability and reproducibility, we incorporate a data-driven hyperparameter tuning step based on Bayesian optimization, enabling users to achieve high-performing and robust hyperparameter configurations with minimal pilot data. We demonstrate the utility of the proposed searchlight optimization framework in an emotion imagination paradigm involving multiple affective states and show that the RSA-based voxel selection approach exhibits complementary characteristics to classifier-based approaches.

## 2. MATERIALS AND METHODS

### 2.1 Experimental Data

The proposed searchlight optimization framework was developed and evaluated using a subset of a neurofeedback dataset from our previous work (Wang et al., 2026).

#### 2.1.1 Design

The study consisted of a single MRI session comprising one anatomical scan and four sequential functional runs. Each functional run followed a task-rest block design, comprising 10 emotion imagery blocks interleaved with 11 baseline blocks, each lasting 20 seconds. Each of the four runs corresponded to one distinct affective state: joyful relaxation, sadness, enthusiasm, or anger. These emotions were selected according to their valence and arousal levels (Feldman, 1995; Heffner & FeldmanHall, 2022). The study was preregistered (https://osf.io/5ybuw) and approved by the Ethics Committee of the University Hospital RWTH Aachen [EK 23-175].

#### 2.1.2 Participants

Six healthy volunteers (4 females, mean age = 26.2 ± 2.5 years) participated in the pilot study, and twenty-seven healthy volunteers (21 females, mean age = 28.7 ± 8.3 years) took part in the main study. The exclusion criteria were acute severe neurological, psychiatric, or somatic disorders. All participants provided written informed consent prior to participation in accordance with the Declaration of Helsinki.

#### 2.1.3 MRI data acquisition

Imaging data was acquired on a 3.0 Tesla Siemens scanner (Siemens Medical Solutions, Erlangen, Germany) with a 20-channel head coil. Anatomical data was obtained using a T1-weighted magnetization-prepared rapid gradient-echo (MPRAGE) sequence (time of repetition (TR): 2000 ms, echo time (TE): 3.03 ms, matrix size: 256×256, voxel size: 1 mm, flip angle: 9°, number of slices: 176, duration: ∼4.3 min). For the subsequent functional runs, a multi-band multi-echo echo-planar imaging (EPI) sequence was used (TR: 1000ms, TE: [13, 27.8, 42.6, 57.4 ms], matrix size: 64×64, voxel size: 3 mm, flip angle: 67°, number of slices: 36, inter-slice gap: 0.75 mm, multi-band factor: 3, GRAPPA factor: 2, partial Fourier phase-encoding factor: 7/8, duration: 7 min).

#### 2.1.4 Data preprocessing

Imaging data was preprocessed in real time using Turbo Brain Voyager (TBV) version 4.2 (Brain Innovation B.V., Maastricht, The Netherlands) and the rt-RSA Python tool (https://github.com/assuntaciarlo/rtRSA_GUI). Specifically, multi-echo images were first combined via a per-volume T2*-weighted sum (Heunis et al., 2021; Mathiak et al., 2002; Weiskopf et al., 2005) in the rt-RSA tool. TBV then performed the preprocessing steps, including motion correction with trilinear interpolation, intra-session alignment to the first volume of the first functional run, and spatial smoothing with a 6 mm full width at half-maximum (FWHM) Gaussian kernel.

The same predefined brain mask as in our previous work (Wang et al., 2026) was used, which comprises emotion-related cortical and subcortical brain regions, with cortical regions defined by the Schaefer atlas (17-network parcellation, 200 parcels, 1 mm resolution) (Schaefer et al., 2018) and subcortical regions defined by the Harvard-Oxford atlas (maximum probability maps, thresholded at 25%, 1 mm resolution) (Desikan et al., 2006).

### 2.2 Searchlight Optimization Framework

The searchlight optimization framework is illustrated in Figure 1. It consists of two main components: a voxel selection step and a hyperparameter optimization step. Pilot data are used to systematically explore and refine hyperparameter configurations, after which the optimized settings are fixed and applied to the main dataset.

**Figure 1.**
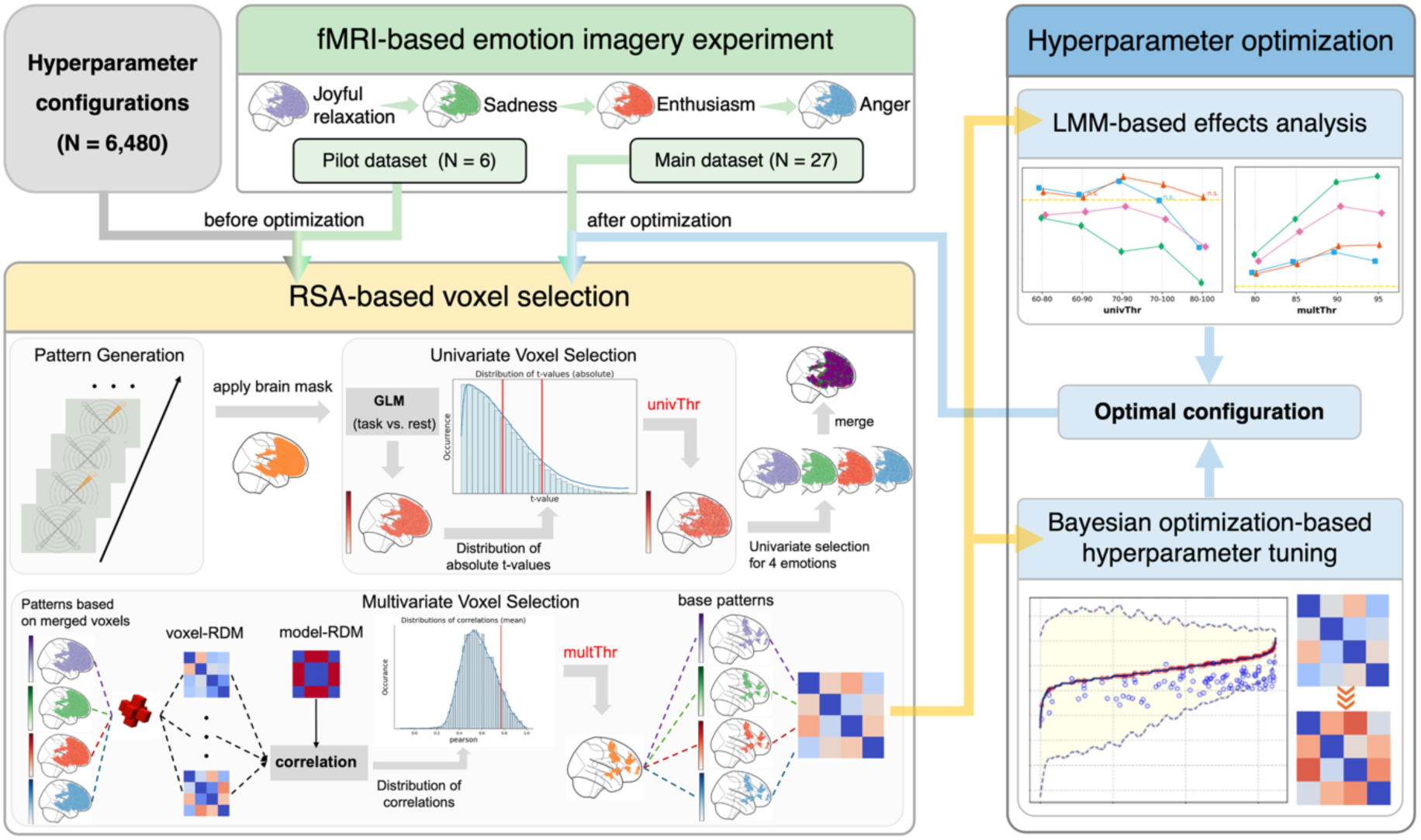
Overview of the searchlight optimization framework. Pilot multi-condition fMRI data are first used to explore voxel selection across a range of hyperparameter configurations. The resulting outputs are then evaluated through hyperparameter optimization, combining linear mixed-effects modeling (LMM) to assess hyperparameter effects and Bayesian optimization to identify optimal configuration. The derived configuration is subsequently fixed and applied to the main dataset. Within the voxel selection approach, functional runs are processed independently through univariate selection and then jointly integrated in a multivariate module to produce the final set of selected voxels.

#### 2.2.1 Voxel selection approach

As shown in Figure 1, the RSA-based voxel selection approach was developed to identify optimal target brain regions that are consistently engaged across multiple mental processes. To initiate the selection, a task-rest block-design functional run for each mental process is required. Optionally beginning with a predefined brain mask, the approach integrates both univariate activation-based and multivariate pattern-based analyses to refine the selection of target regions. The approach was primarily implemented in the rt-RSA tool, with statistical modelling conducted in TBV.

##### Univariate selection

The univariate selection was performed using a voxel-wise general linear model (GLM) applied separately to each functional run. For each voxel within the predefined brain mask, t-values were computed for the contrast ‘task versus rest’, generating a statistical activation map per run (i.e., per condition). Voxels were then ranked according to the absolute t-value distribution, and those within a specified percentage interval (defined by the hyperparameter *univThr*, e.g., 60-90%) were selected as candidates for the univariate selection results. To minimize the influence of small or isolated activations (Geerligs & Maris, 2021), clusters containing fewer voxels than the threshold (defined by the hyperparameter *clusterThr*) were excluded.

This procedure was repeated independently for each functional run, producing a binary selection map per run. The final univariate selection map was obtained by combining all run-specific selection maps using a logical OR operation, ensuring the inclusion of voxels that were strongly activated in at least one run.

##### Multivariate selection

The multivariate selection was performed using voxel-wise RSA within a searchlight analysis. The univariate selection map was first applied to each functional run to extract the t-values of the corresponding voxels, resulting in multiple run-specific univariate patterns. After specifying the searchlight *shape* and *radius* hyperparameters, voxel-wise representational dissimilarity matrices (RDMs) were computed. Specifically for each voxel, a local pattern vector was defined by the t-values of all voxels within its searchlight (centered at that voxel), and pairwise Pearson’s correlations between run-specific local pattern vectors were calculated to construct the voxel-level RDM.

Each voxel’s searchlight-based representational structure (i.e., the searchlight-based voxel RDM) was then compared with a target model RDM using a similarity metric (Pearson’s or Spearman correlation, defined by the hyperparameter *metric*). The target model RDM encoded the hypothesized similarity structure among task conditions and could be flexibly defined to reflect different representational assumptions, such as categorical or graded condition differences (Figure 2). The resulting similarity map was spatially smoothed within each searchlight according to the hyperparameter *mode*, which determined whether the final voxel similarity value corresponded to the voxel itself, or to the mean or median of its local neighborhood. Voxels within the top percentage interval of the similarity distribution (defined by the hyperparameter *multThr*) were retained as candidates for the multivariate selection results. Similar to the univariate selection stage, small clusters containing fewer voxels than *clusterThr* were iteratively removed until no clusters could be excluded or the remaining voxel count reached the lower limit (defined by the hyperparameter *voxelThr*). The remaining voxels formed the final multivariate selection map.

**Figure 2.**
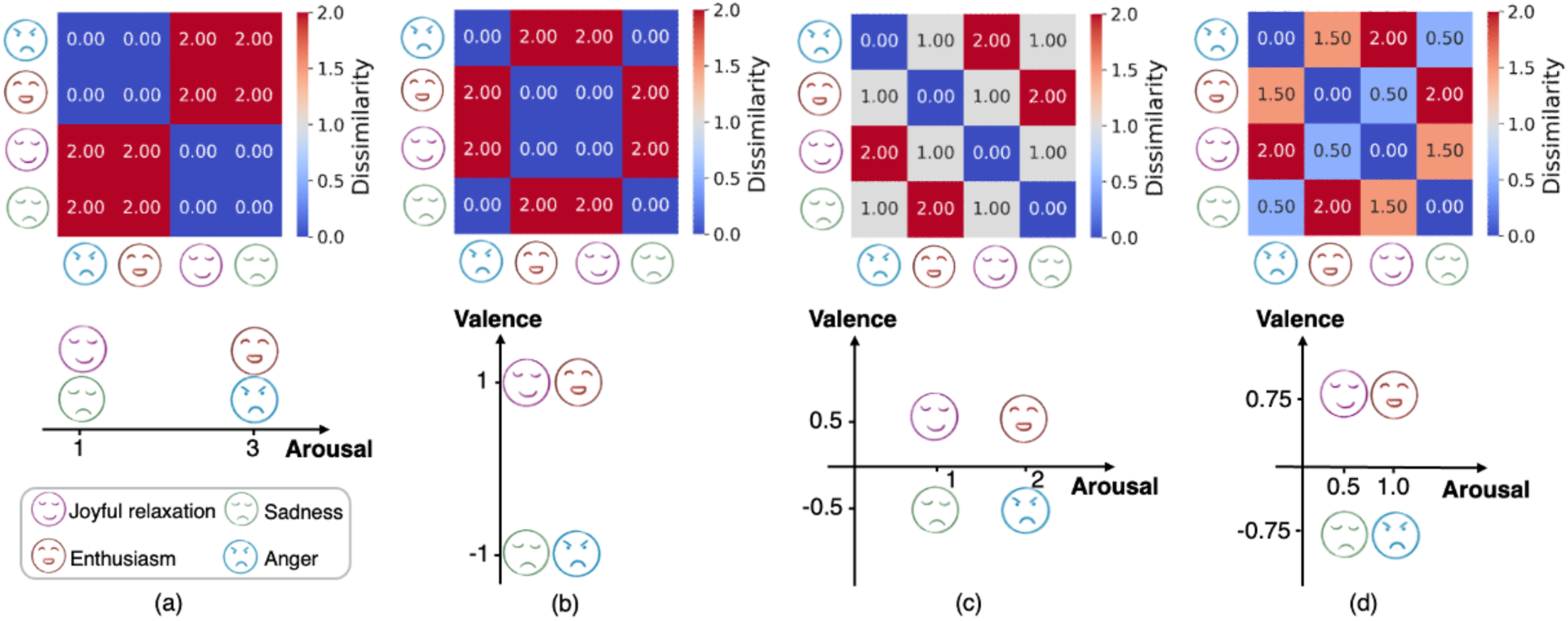
Target model RDMs. All RDMs were defined in a valence–arousal representational space, with pairwise dissimilarities computed using Manhattan distance (maximum = 2). Panels illustrate different modeling assumptions: (a) arousal-only model, encoding variation exclusively along the arousal dimension; (b) valence-only model, capturing differences between positive and negative emotions; (c) combined model with equal weighting of valence and arousal; and (d) combined model with unequal weighting of the two dimensions.

For performance evaluation, t-values of voxels in the multivariate selection map were extracted from each functional run and used as run-specific multivariate patterns. The relationships among these multivariate patterns were assessed by an empirical RDM, computed from their pairwise dissimilarities. Performance was then quantified as the Spearman correlation between this empirical RDM and the target model RDM (Kriegeskorte et al., 2008; Nili et al., 2014), reflecting the degree to which the selected voxels preserved the hypothesized representational structure.

#### 2.2.2 Hyperparameter optimization

To enhance the practical feasibility of the RSA-based voxel selection approach, we incorporated an efficient hyperparameter optimization step and evaluated multiple target representational patterns with varying levels of complexity. Specifically, to determine an optimal configuration, a range of plausible values were systematically tested for each hyperparameter, with consideration of both computational efficiency and practical applicability (Table 1). In addition, to assess the impact of different representational targets, four model RDMs were defined along the valence and arousal dimensions, as illustrated in Figure 2. Two models captured single affective dimensions, whereas the other two combined both dimensions. Notably, the final model introduced graded dissimilarity values by weighting the two dimensions differently, in contrast to the binary values used in the other three models, thereby increasing representational complexity. Including these models allows us to examine how the searchlight-based selection performs when incorporating additional affective dimensions and varying levels of representational complexity. Method performance was evaluated using the Spearman correlation between the final selected voxels’ RDM and the target model RDM.

**Table 1.** Hyperparameters and corresponding levels.

| Hyperparameters | Definition | Levels |
| --- | --- | --- |
| *shape* | Searchlight shape | sphere, cube |
| *radius* | Searchlight radius | 1, 2 |
| *metric* | Method to correlate model RDM and voxel RDM | Pearson, Spearman |
| *mode* | How to compute voxel RDM based on searchlight | center, mean, median |
| *univThr* | Threshold to filter voxels based on absolute t-values | 50-80, 60-90, 70-100,<br>60-80, 70-90, 80-100 |
| *multThr* | Threshold to filter voxels based on correlations | 75, 80, 85, 90, 95 |
| *clusterThr* | the minimum size for a cluster | 10, 15, 20 |
| *voxelThr* | the minimum number of selected voxels | 100, 200, 300 |

##### Hyperparameter effects on performance

We examined how the hyperparameters influenced overall performance by treating each hyperparameter level as a categorical factor. A linear mixed-effects model (LMM) was used to assess main effects on performance, with participants included as a random effect to account for inter-individual variability. To quantify the variability, the intraclass correlation coefficient (ICC) was calculated based on the variance components estimated from the LMM. Multiple comparisons across parameter levels were controlled using false discovery rate (FDR) correction, applied separately for each hyperparameter.

##### Hyperparameter tuning

Based on the parameter ranges in Table 1, a total of 6,480 hyperparameter combinations were defined. Using the pilot dataset (N = 6), we first conducted a grid search across all combinations to obtain the complete performance landscape. Bayesian optimization was then performed on the same dataset for 100 iterations to identify an approximate optimal configuration. Specifically, following previous Bayesian optimization work (Shahriari et al., 2016; Snoek et al., 2012), the objective function (Spearman correlation) was modeled using a Gaussian process surrogate with a Matern kernel (ν = 2.5), as implemented in the scikit-optimize python module (Head et al., 2021). At each iteration, an acquisition function (expected improvement) was used to balance exploration and exploitation when selecting candidate configurations. The resulting configurations were evaluated and positioned within the full performance landscape derived from the grid search. To further evaluate the generalizability of the fine-tuned configurations, the same 100 combinations obtained from Bayesian optimization on the pilot dataset were directly applied to the main dataset (N = 27). Performance measures from both datasets were then compared using Pearson’s correlation.

### 2.3 Performance analysis

The optimal hyperparameter configuration for subsequent performance analyses was determined by running Bayesian optimization with ten different random seeds and selecting the most frequently occurring configuration among the top ten candidates. The hyperparameter tuning step was conducted independently for each target model RDM. The resulting optimal configurations were then used to evaluate performance improvements across selection stages and to compare the RSA-based voxel selection approach with the classifier-based approach.

#### Improvement across internal stages

As shown in Figure 1, the RSA-based voxel selection approach produced selection results at two stages: (i) voxels retained after univariate selection, and (ii) voxels retained after multivariate selection. To examine performance variations between them, we calculated the final RDMs derived from the corresponding selected voxels across the four conditions.

#### Comparison to classifier-based approach

For comparison, we replaced the RSA-based voxel selection approach in our framework with a classifier-based approach using the same predefined brain mask as starting point. All functional data and motion parameters were taken from the real-time TBV output, identical to the preprocessing pipeline used in the RSA-based approach.

Classification samples were generated using block-wise GLM analyses, in which each pair of consecutive task and rest blocks was modeled separately to obtain task-vs-rest t-maps. This windowed procedure yielded one sample per block pair, resulting in 40 samples across the four runs (10 per emotion). The GLM design matrices included standard motion confounds derived from TBV real-time output. The samples were then grouped according to the classification scheme corresponding to each target model RDM: binary classification for single-dimension models and four-class classification for combined valence–arousal models. Voxel-wise classification features were defined using a searchlight framework that matched the searchlight shape and radius determined by the fine-tuned hyperparameters of the RSA-based approach. Within each searchlight, logistic regression (LR) with L2 regularization was applied using five-fold cross-validation across the 40 samples. The resulting mean classification accuracy for each searchlight was assigned to its center voxel, generating a voxel-wise classification-accuracy map.

To obtain a final voxel set, voxels were selected based on classification accuracy to match the number of voxels in the RSA-based approach. For comparison with our selection results, spatial differences between the two resulting voxel sets were visualized on brain maps. In addition, the corresponding t-values from the full-run t-maps were extracted for both voxel sets to compute RDMs and whole-run classification accuracies. Statistical differences between approaches were assessed using paired t-tests across participants.

## 3. RESULTS

### 3.1 Hyperparameter effects on performance

Results of the LMM analysis assessing the main effects of all hyperparameters on overall performance are shown in Table 2. Overall, most hyperparameters showed significant effects on performance. For the searchlight configuration, performance estimates were higher for the cube-shaped searchlight compared with the spherical one, and a larger searchlight radius (2 voxels) produced higher estimates relative to the reference radius (1 voxel). Regarding (dis)similarity computation, Pearson’s correlation was linked with increased performance compared to Spearman correlation, and incorporating information from the entire searchlight neighborhood (using mean or median value) showed higher estimates than relying only on the central voxel. Among amplitude-related parameters, cluster size and total voxel count showed relatively small effects on performance. To facilitate interpretation of hyperparameters with multiple levels, the effects of the two selection thresholds are additionally visualized in Figure 3. Consistent with the estimates reported in Table 2, the univariate threshold showed relatively stable performance across lower-to-mid percentile ranges, followed by a decrease at higher thresholds. In contrast, the multivariate threshold was associated with increasing performance at lower percentile ranges, with effects stabilizing at higher levels.

**Figure 3.**
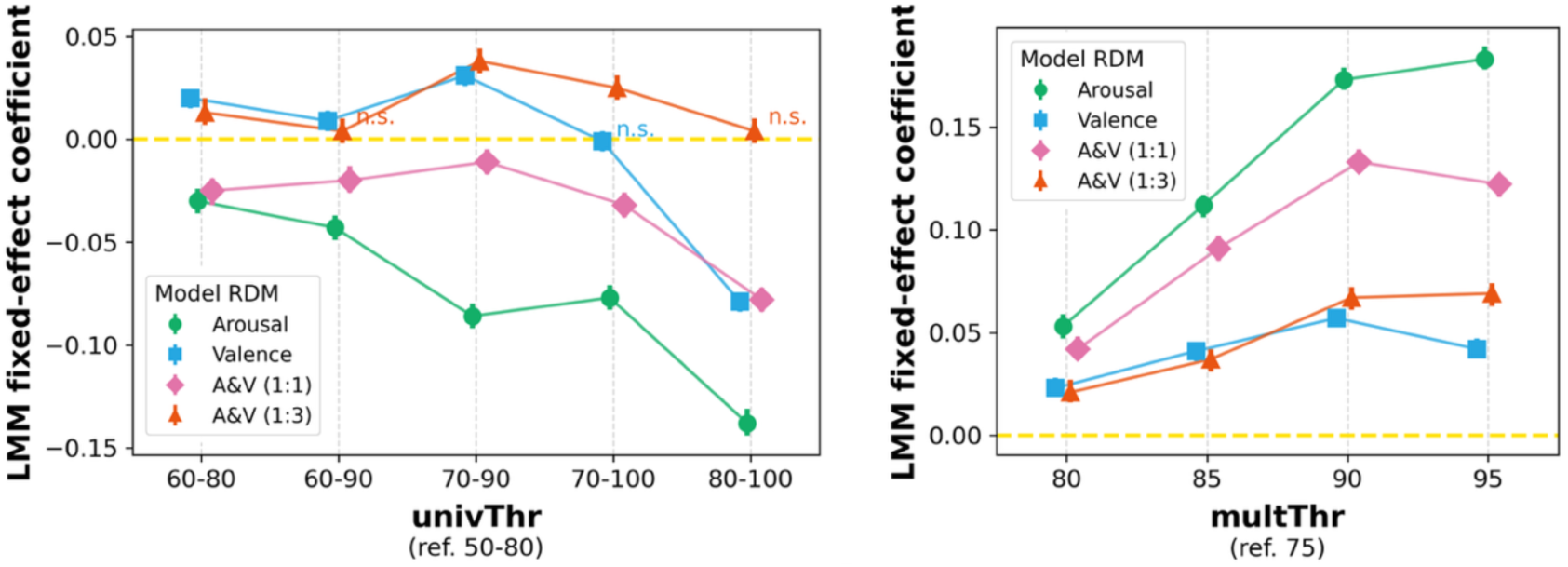
Main effects of the two selection thresholds (univariate and multivariate) from the LMM analysis. Each panel displays the estimated fixed-effect coefficients and their 95% confidence intervals for different parameter values relative to the reference value (yellow line), which is indicated under each hyperparameter term. Different colors within each panel correspond to results derived from different target model RDMs. Non-significant effects (p ≥ 0.05) are labelled as “n.s.”.

**Table 2.**
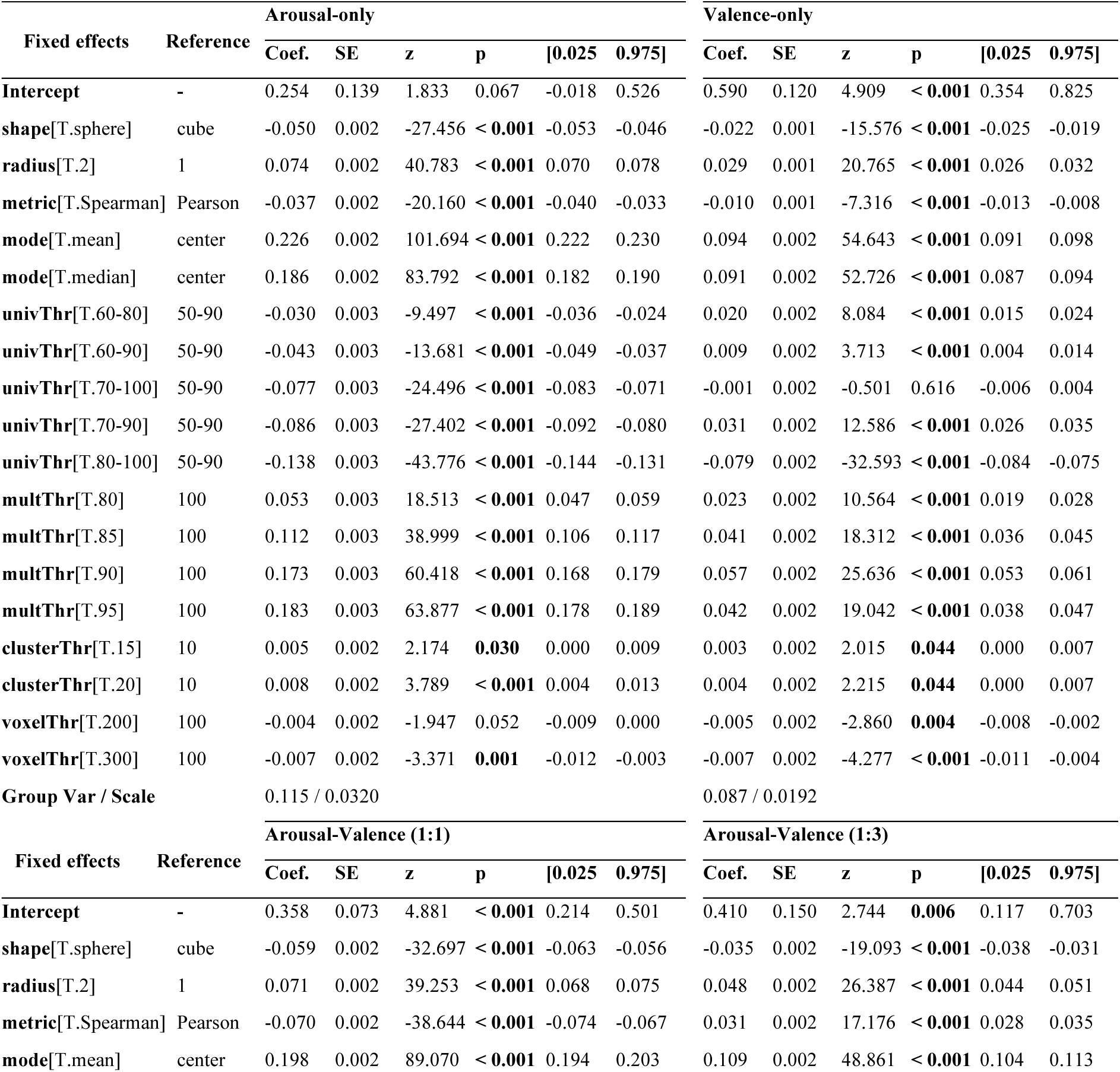
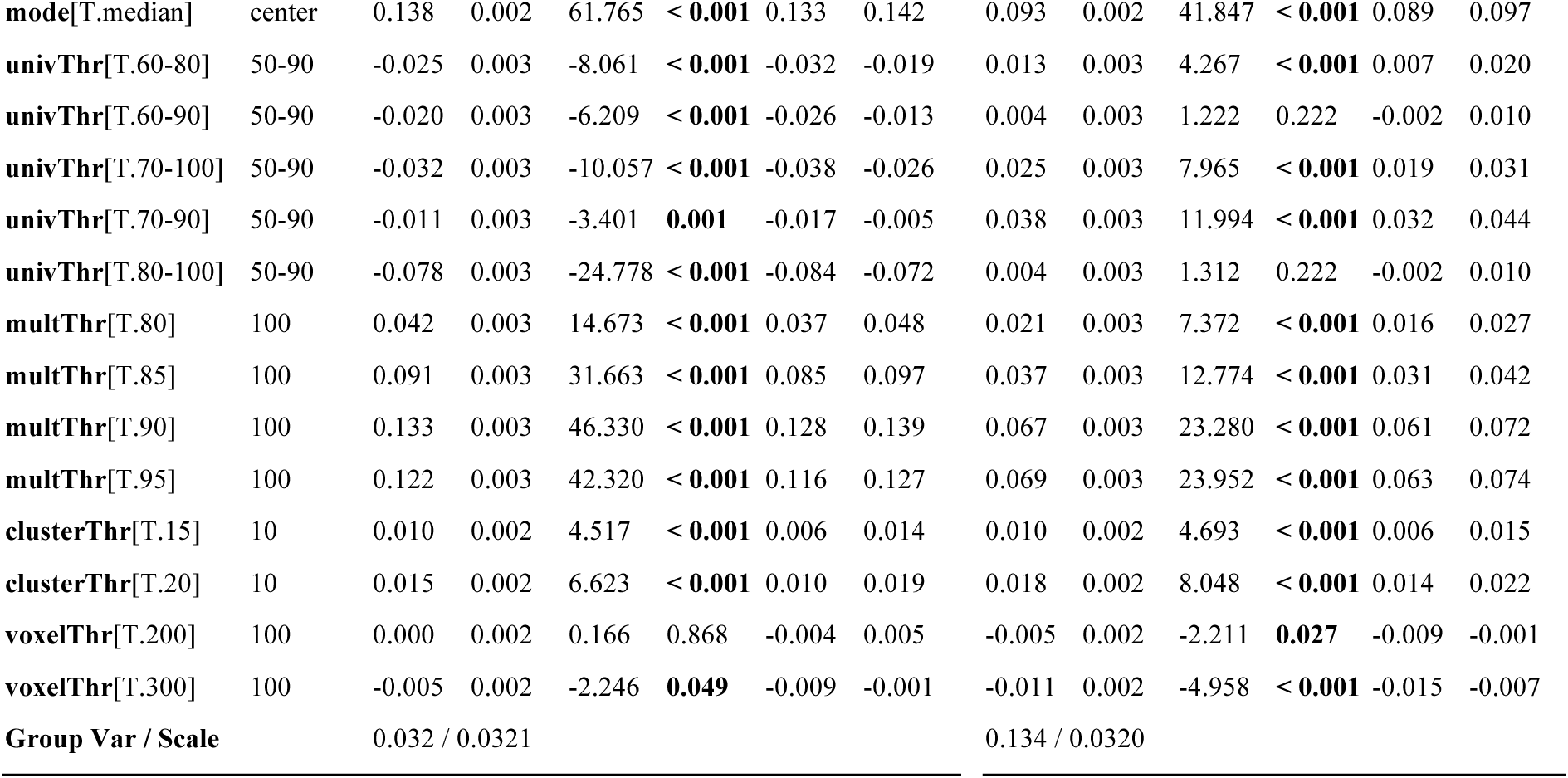
Main effects from the LMM analysis (6 participants, 6480 observations per participant). For each hyperparameter, estimated fixed-effect coefficients (Coef.), standard errors (SE), z-statistics (z), p-values (p), and 95% confidence intervals ([0.025, 0.975]) are reported relative to the reference level. Variance components, including variance across individuals (Group Var) and residual variance (Scale), are also provided. P-values were corrected for multiple comparisons using FDR within each <u>hyperparameter group, except for parameters with only two lev</u>els, for which no correction was applied.

In addition, most hyperparameters exhibited consistent effect patterns (Table 2), and the *univThr* and *multThr* selection thresholds showed similar trajectories across parameter ranges and target model RDMs (Figure 3). These observations indicate that the proposed optimization framework generalizes across model specifications. Differences were primarily observed when rank complexity increased from a binary to a graded structure (unequal combined model). For instance, in the more complex arousal-valence model (red line in Figure 3), varying the univariate threshold did not significantly influence performance, whereas it remained a key factor in the other models (Figure 3). Furthermore, under this complex graded model, Spearman correlation outperformed Pearson’s correlation (Table 2), whereas Pearson’s correlation showed higher estimates in the other models, which may indicate that the neural data aligned more closely with the ordinal structure of the target model than with its exact linear spacing.

Lastly, variance components indicated moderate to substantial inter-subject variability (ICC range: 0.50–0.82), while the narrow confidence intervals (CIs) reflect high precision driven by the large number of within-subject observations (i.e., hyperparameters included in the analysis). In contrast, the confidence interval estimates around the intercept (which is based on only a combination of the reference values for all hyperparameters) are consistently larger across all four RDMs. Overall, this suggests relatively stable within-subject hyperparameter effects, although results should be interpreted cautiously given the small number of participants (N = 6).

### 3.2 Efficient hyperparameter tuning

The grid search results across all 6,480 parameter combinations on the pilot dataset (N = 6) were ordered by Spearman correlation between actual and target RDMs for visualization (independently for each target model RDM; Figure 4). The 100 hyperparameter configurations identified through Bayesian optimization on the pilot dataset (N = 6; red circles in Figure 4) clustered near the global performance maximum within the full grid-search landscape. When these same configurations were directly applied to the main dataset (N = 27; blue circles in Figure 4), their performance showed significant pilot-main correspondence. Besides the Pearson’s correlations shown in Figure 4, Spearman correlations were also significant (p < 0.001) across all target models (ρ = 0.755, 0.603, 0.556, and 0.507 for panels (a)–(d), respectively), indicating a consistent relative ranking of hyperparameter sets across samples. These findings demonstrate that Bayesian optimization can efficiently approximate high-performing hyperparameter configurations using a small pilot sample, which can then be generalized to larger datasets.

**Figure 4.**
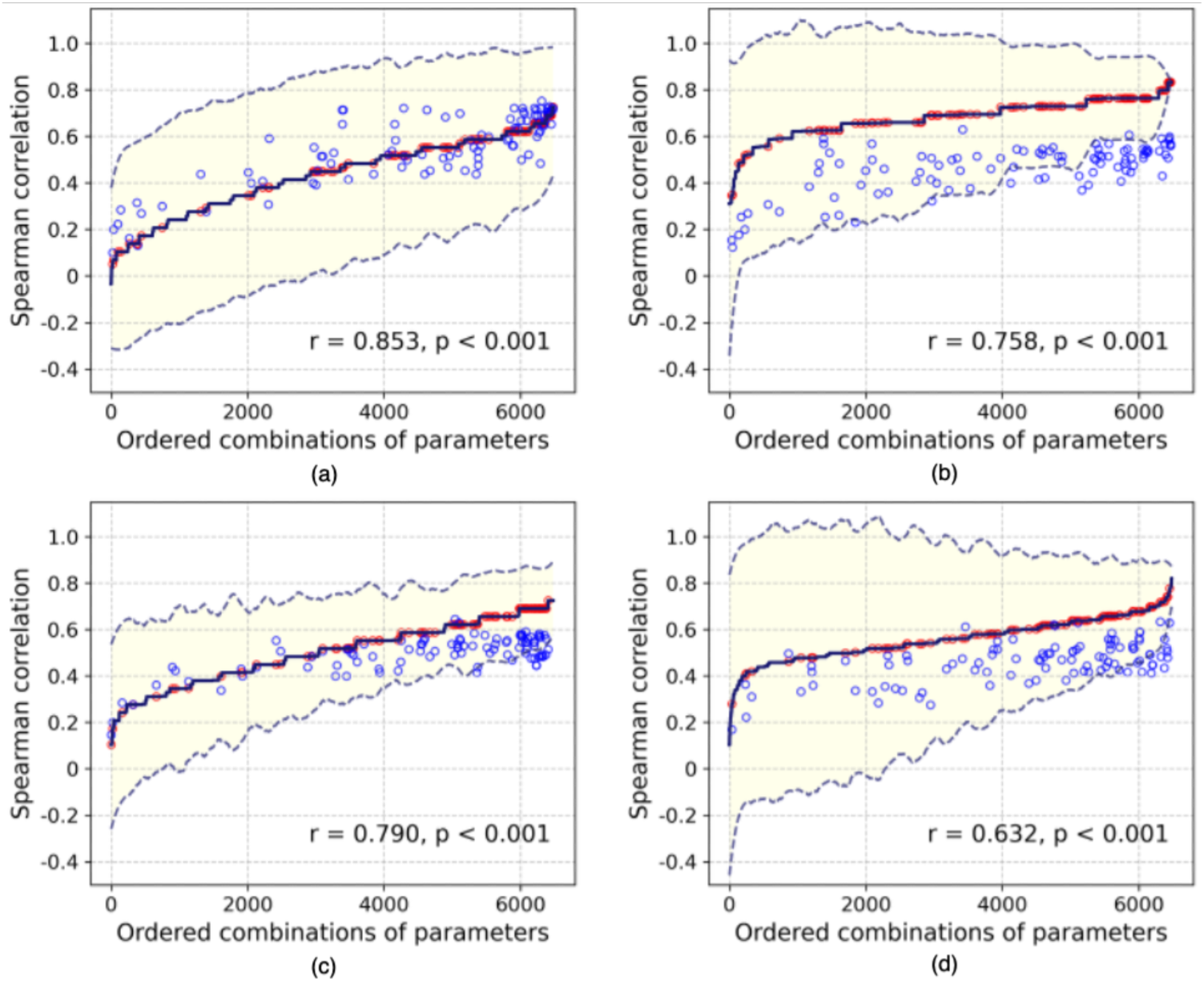
Hyperparameter search and generalization performance. The solid curve and shaded area represent the grid search results across all 6,480 hyperparameter combinations on the pilot dataset (mean ± 95% CI across six participants). Red circles indicate the 100 Bayesian optimization trials performed on the pilot dataset, while blue circles show the performance of the same 100 hyperparameter configurations when directly applied to the main dataset (N = 27). The Pearson’s correlation between pilot and main dataset performance is reported at the bottom of each panel. Panels correspond to different target model RDMs: (a) arousal-only, (b) valence-only, (c) arousal–valence (1:1), and (d) arousal–valence (1:3).

Target model-specific differences were observed alongside the overall consistent tuning effectiveness, where distinct patterns emerged between the arousal-only and valence-only models. For the arousal-only model (Figure 4(a)), mean performance in the main dataset (N = 27) was comparable to or slightly higher than that in the pilot dataset (N = 6), whereas for the valence-only model (Figure 4(b)), performance was consistently lower in the main dataset. This pattern may suggest a relatively stable arousal-related representational structure across participants in the present task. In addition, the CIs narrowed with improved hyperparameter settings for the valence-only model (Figure 4(c)) but remained constant for the arousal-only model (Figure 4(a)). The stable CI width in the arousal-only model suggests relatively consistent between-participant variability across parameter configurations, whereas the narrowing pattern in the valence-only model indicates reduced variability under better-performing settings. The combined models (Figures 4(c) and 4(d)) exhibited intermediate patterns consistent with weighted contributions from the two single-dimension models.

### 3.3 Performance analysis

The optimal hyperparameter configurations used for subsequent performance analyses are summarized in Table 3. Based on these fixed settings, we evaluated performance changes across its internal stages and compared it with an external classifier-based voxel selection approach.

**Table 3.** Optimal hyperparameter configurations determined by repeated Bayesian optimization for each target model RDM.

| Parameters | Arousal-only | Valence-only | Arousal-Valence (1:1) | Arousal-Valence (1:3) |
| --- | --- | --- | --- | --- |
| shape | sphere | cube | cube | cube |
| radius | 2 | 1 | 2 | 2 |
| metric | Spearman | Pearson | Pearson | Spearman |
| mode | mean | mean | mean | mean |
| univThr | 50-80 | 60-80 | 50-80 | 60-80 |
| multThr | 95 | 85 | 85 | 95 |
| clusterThr | 10 | 20 | 10 | 15 |
| voxelThr | 300 | 100 | 300 | 100 |

#### 3.3.1 Improvement across internal stages

Internal performance was assessed by comparing the RDMs derived from voxels selected at the univariate and multivariate stages of the RSA-based voxel selection approach (Figure 5). After univariate selection, the resulting RDM showed dissimilarities (one minus Pearson’s correlation) around 0.5 across pairwise conditions, indicating limited discernibility among the four emotional conditions. Some univariate-stage RDMs were identical or nearly identical across target models, which was expected because univariate selection depends only on the optimized *univThr* and *clusterThr* parameters. Consistent with the optimal configurations in Table 3, models sharing the same or highly similar values for these parameters therefore produced the same or nearly identical voxel sets. In contrast, the RDM following multivariate selection showed greater similarity to the corresponding target model RDM (Figure 2), demonstrating improved separability between emotional conditions and stronger alignment with the hypothesized representational structure among these affective states.

**Figure 5.**
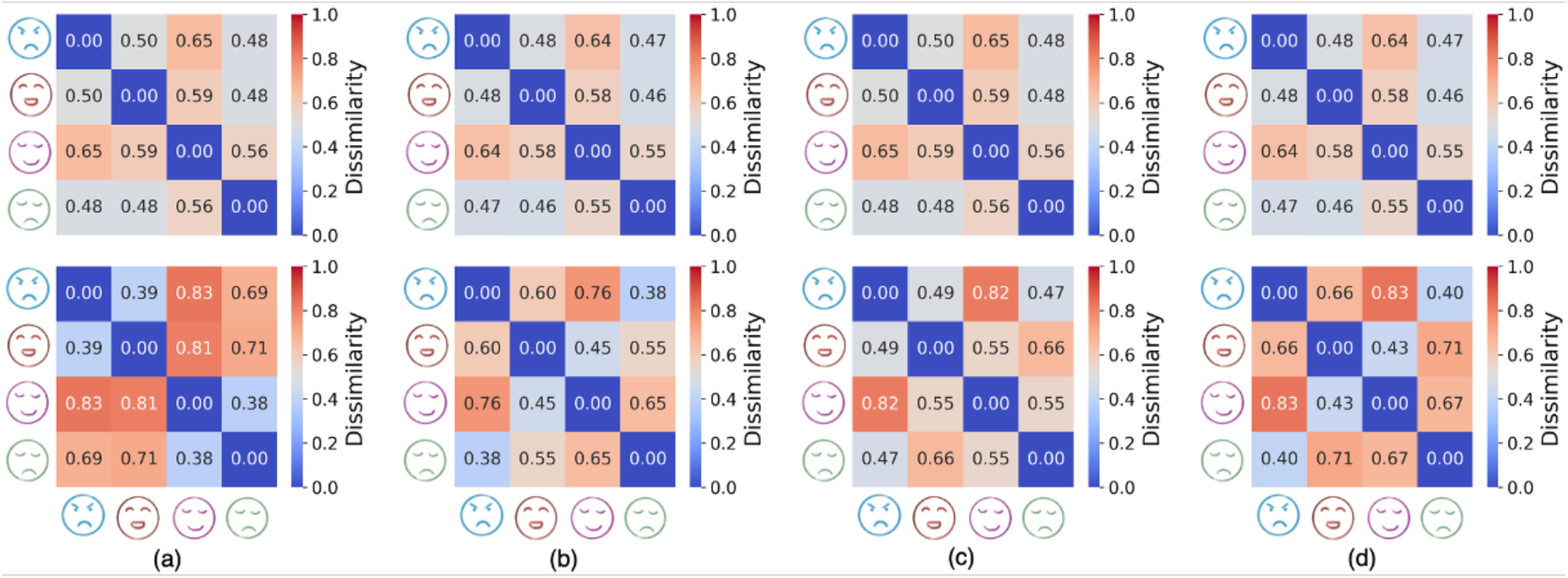
RDMs derived from univariate and multivariate selection stages. Each matrix represents the group-level averaged RDM computed from voxels selected after univariate (top row) and multivariate (bottom row) stages. Columns correspond to target model RDMs: (a) arousal-only, (b) valence-only, (c) arousal–valence (1:1), and (d) arousal–valence (1:3).

#### 3.3.2 Comparison with classifier-based approach

Figure 6 illustrates both the spatial overlap and differences between voxels selected by the RSA-based and LR-based approaches. In Figure 6(a), the two approaches showed substantial overlap across target models, with Dice coefficients of 0.652 for the arousal-only model, 0.848 for the valence-only model, 0.910 for the 1:1 arousal–valence model, and 0.583 for the 1:3 arousal–valence model. This overlap was mainly observed in orbitofrontal, broader prefrontal, and limbic regions previously identified as being involved in emotion imagery (Linhartová et al., 2019), supporting the general effectiveness of both approaches in identifying emotion-related voxels. Figure 6(b) further illustrates the difference in voxel selection frequency across participants. Regions with values near zero indicate comparable selection frequency between approaches, whereas positive and negative values reflect relative dominance of the RSA-based and LR-based approaches, respectively. The LR-based approach showed relatively greater selection in lateral prefrontal regions, including inferior and middle frontal gyri. In contrast, the RSA-based approach more frequently selected medial prefrontal and limbic regions, particularly the anterior cingulate cortex (ACC), amygdala, and hippocampus. These differences were more pronounced in the combined models, where both arousal and valence dimensions were incorporated. Overall, while both approaches captured core emotion-related regions, they exhibited partially distinct spatial preferences across prefrontal and limbic areas.

**Figure 6.**
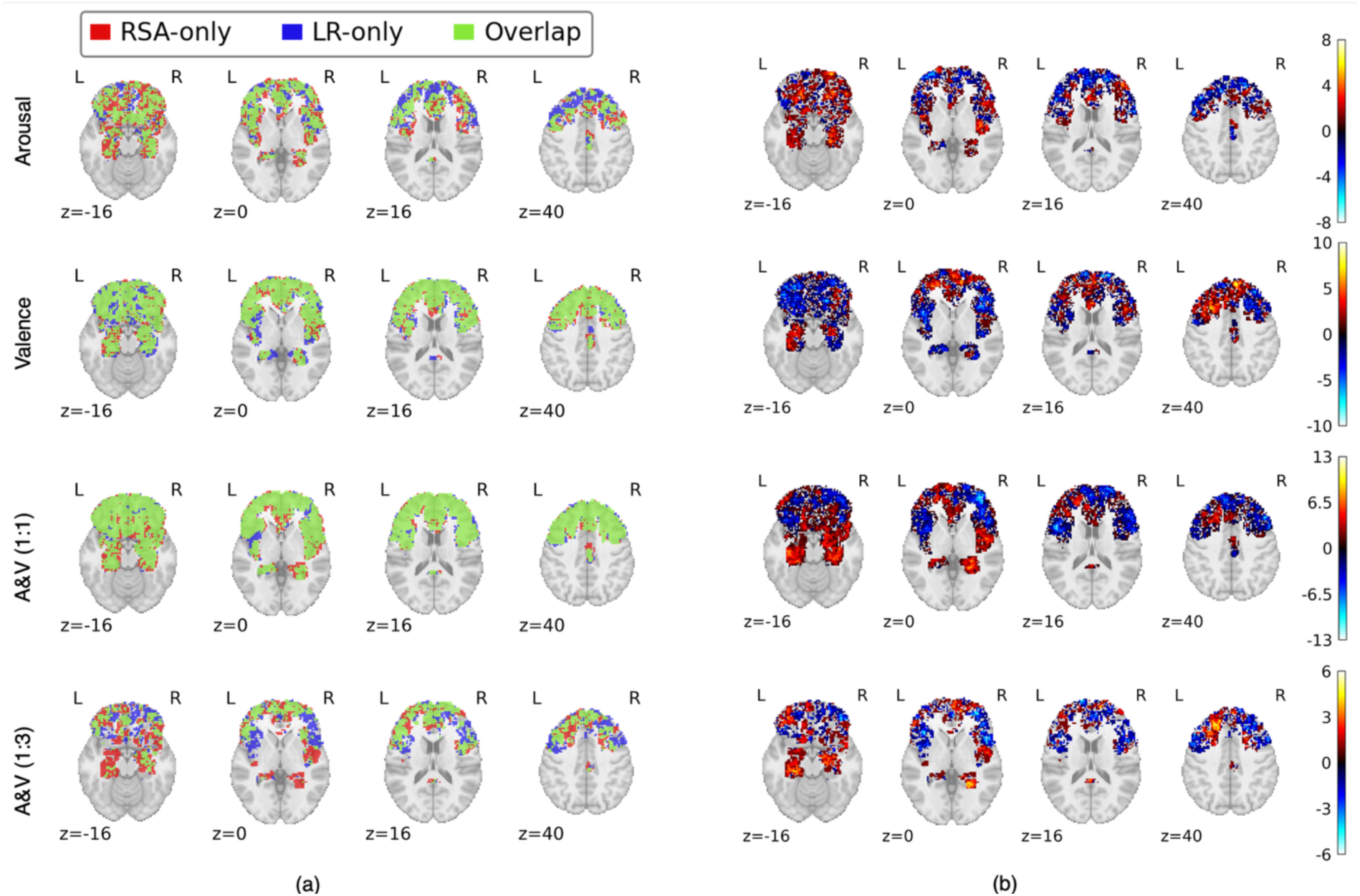
Spatial comparison of voxels selected by the RSA-based and LR-based approaches. (a) Overlap maps, with red, blue, and green indicating RSA-only, LR-only, and overlapping voxels, respectively. (b) Differences in selection frequency across 27 participants, computed as RSA minus LR; positive and negative values indicate more frequent selection by the RSA-based and LR-based approaches, respectively. Rows correspond to target model RDMs.

Using Spearman correlation between empirical and target RDMs (Figure 7(a)) and classification accuracy (Figure 7(b)) as complementary performance metrics, we compared voxels selected by the RSA-based and LR-based approaches. When performance was assessed using Spearman correlation, the RSA-based approach yielded significantly higher values than the LR-based approach across all target models (all p < 0.001). Although the LR-based approach produced positive correlation values, statistical significance was observed only for the arousal-only model (p = 0.005), with the remaining models failing to reach significance (p > 0.05). In contrast, when evaluated by classification accuracy, the LR-based approach outperformed the RSA-based approach across all target models (all p < 0.001). Nevertheless, classification accuracy of the RSA-based voxels remained significantly above chance level for all models (all p < 0.001), indicating that these voxels retained reliable discriminative information.

**Figure 7.**
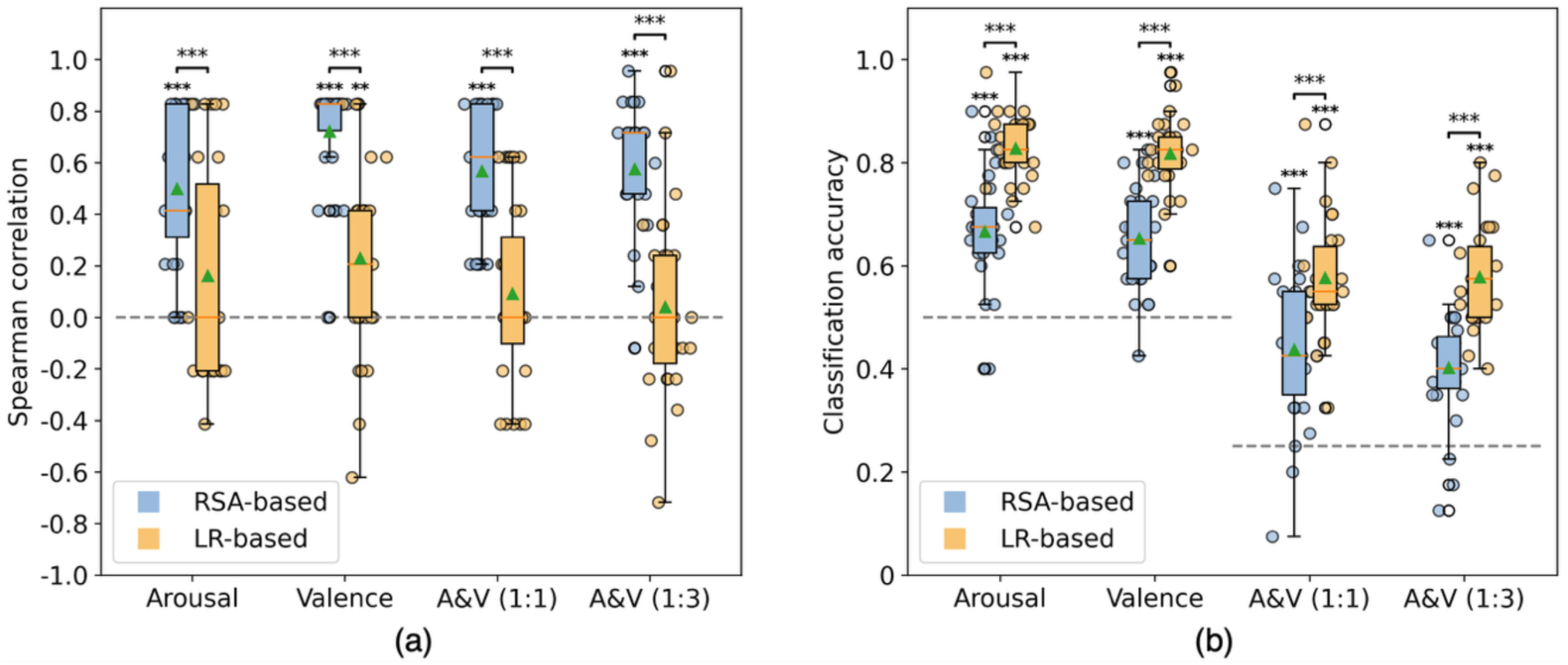
Representational and classification performance of the RSA-based and LR-based voxel selection approaches. (a) Spearman correlations between individual empirical RDMs and the corresponding target model RDMs; the gray line indicates the null value (ρ = 0). (b) Classification accuracies across participants; gray lines indicate chance levels of 0.50 for single-dimension models (arousal-only and valence-only) and 0.25 for combined arousal–valence models (arousal–valence 1:1 and 1:3). Boxplots show the median, IQR, 1.5 × IQR whiskers, outliers, and mean (green triangle). Asterisks above individual boxes indicate comparisons against the null/chance level, whereas asterisks above paired boxes indicate comparisons between approaches (*p < .05, **p < .01, ***p < .001).

## 4. DISCUSSION

This study introduces a framework to identify informative voxels for multivariate fMRI studies in a systematic way using pilot data. The framework allows exploring a large hyperparameter space and assessing potential performance gains across target models, as well as their generalizability to an independent larger dataset. Specifically, we propose a subject-level searchlight optimization framework for systematic voxel selection in multi-condition fMRI paradigms and evaluate its performance using a real-time fMRI emotion-imagery dataset. The results provide three main insights. First, the proposed framework integrates voxel-wise GLM-based univariate activation analysis with searchlight-wise RSA-based multivariate pattern analysis, enabling the identification of voxels that are both functionally engaged and capable of distinguishing among related mental conditions. Second, we demonstrate a data-driven hyperparameter tuning step based on Bayesian optimization, efficiently identifying high-performing configurations which show consistent performance from pilot to larger samples. Third, internal between-stage validation and comparison with a classifier-based selection approach indicate that the RSA-based approach preserves structured representational relationships while maintaining discriminative capacity. The framework validity is supported by the convergence of selected voxels with established emotion-related brain regions and by improved alignment with target representational structures. Together, these findings highlight the methodological effectiveness and practical utility of the searchlight optimization framework in addressing key challenges in biomedical signal engineering associated with high-dimensional neuroimaging data. Although evaluated in the context of a multi-condition cognitive and affective fMRI paradigms, the framework can be adapted and extended to other task domains, RSA methodologies, and neuroimaging modalities for studying representational geometry.

The proposed searchlight optimization framework addresses two key challenges in the task of target voxel/region localization for multi-condition fMRI paradigms. First, many previous studies rely on anatomically predefined ROIs or simple GLM-based localizers to define target voxels (Fede et al., 2020; Tschentscher et al., 2024; Tursic et al., 2020; Y. Zhang et al., 2025). While effective for isolating broad or well-localized functions (Zhao et al., 2023; Zotev et al., 2024), such approaches may be less suitable when finer-grained differentiation among closely related conditions is required (Hennings et al., 2022; Valente et al., 2019). By incorporating multivariate information on top of univariate activation results, the proposed framework refines voxel selection to identify regions that are both functionally engaged and sensitive to structured differences across conditions. Second, although classifier-based searchlight decoding is a widely used approach for integrating multivariate information (Kriegeskorte et al., 2006), it requires classifier training and cross-validation at each searchlight location, resulting in substantial computational demands (Bressler et al., 2024; Etzel et al., 2013). In contrast, the RSA-based computation used here relies instead on correlation-based comparisons combined with thresholding and clustering (Weaverdyck et al., 2020), enabling whole-brain processing within seconds. Together, the proposed framework provides an effective and computationally efficient solution for voxel selection in studies requiring differentiation across related mental states.

Another contribution of this work is the introduction and validation of a systematic data-driven hyperparameter tuning step using Bayesian optimization. Voxel selection approaches often involve several predefined hyperparameters, such as threshold ranges and searchlight specifications, which are often determined based on prior experience (Etzel et al., 2013; Jolly & Chang, 2021). Such choices may fail to achieve optimal selection performance and consequently require larger sample size than necessary (Mumford, 2012). The Bayesian optimization-based hyperparameter tuning step implemented here identifies high-performing hyperparameter combinations with only a small number of iterations in a small pilot data set, which overall showed a maintained trajectory of performance gain in a larger sample (Figure 4). However, we also note that the shape of performance trajectories flattened for the two valence-dominant models compared to the arousal-dominant models. This observation is in line with previous multivariate fMRI studies that have shown higher discriminability and generalizability to discriminate low vs. high arousal states compared to or across different valence states (Viinikainen et al., 2010; White et al., 2026; R. Zhang et al., 2025). Moreover, it is in line with the general observation that small pilot data sets tend to overestimate effect sizes compared to larger replications, and they should hence not be used as the sole basis for a sampling plan of a new study(Albers & Lakens, 2018; Lakens, 2022). However, crucially, the presented framework provides trajectories of effect sizes as a function of hyperparameter choices and shows that the rank of these trajectories was largely preserved as shown by the generally high correlations between pilot and main data set performance (Figure 4). On this basis, these trajectories provide more informative, stable insights for sample size planning compared to conventional single point effect size estimates. Overall, the systematic hyperparameter tuning step therefore reduces dependence on arbitrary hyperparameter choices and provides a practical route toward high-performance voxel selection. It thereby allows drawing more information from pilot data, which can be used when planning and pre-registering a new study (Allen & Mehler, 2019).

Importantly, because statistical power in correlation-based analyses depends strongly on effect size, improvements in representational alignment directly translate into reduced sample size requirements. For example, increasing the representational correlation from a moderate effect (e.g., ρ ≈ 0.35) to a stronger effect (e.g., ρ ≈ 0.60) would reduce the required sample size for 90% power (two-tailed) from approximately 78 participants to about 19, representing a reduction of more than 70%. Although the exact magnitude depends on model specification and representational complexity, it illustrates how principled hyperparameter tuning can enhance statistical efficiency and improve the feasibility of multi-condition fMRI research (Algermissen & Mehler, 2018). Moreover, we note that this gain in power will potentially increase for more complex hypotheses and corresponding models that are tested. Such gains in statistical sensitivity become also in particular relevant for studies that either directly target (Ciarlo et al., 2022; Russo et al., 2021; Wang et al., 2026) or assess (Kikumoto et al., 2025) changes in representational geometry as outcome measures following an intervention.

In addition, a noteworthy observation from the hyperparameter tuning results relates to the univariate threshold. Conventional GLM-based voxel selection approaches typically retain only the strongest-activated voxels by applying a top-percentile threshold (Lee et al., 2024), under the assumption that the highest task-related activations carry the most informative signal. Interestingly, our fine-tuned hyperparameter configurations consistently favored middle percentile ranges (e.g., 50–80%) rather than the highest activations. This finding suggests that voxels with moderate but reliable task-related activation may be more effective for distinguishing among related conditions (Tarhan & Konkle, 2020; Walther et al., 2016), further highlighting the importance of integrating univariate and multivariate information when selecting voxels for multi-condition paradigms.

Applying the fine-tuned hyperparameters to the main emotion-imagery dataset further supported the effectiveness of the proposed framework. Comparison of RDMs derived from voxels selected at the univariate and multivariate stages revealed a clear refinement effect: the multivariate selection produced voxel sets with substantially stronger alignment to the corresponding target model RDMs across all tested models. In contrast, RDMs obtained after univariate selection showed weaker representational alignment. These findings indicate that incorporating multivariate representational information enhances sensitivity to structured differences among conditions beyond activation-based selection alone.

Moreover, the comparison with the classifier-based voxel selection approach highlights the complementary strengths of the two approaches. As expected, each approach performed best on the metric it was implicitly optimized for: the classifier-based approach achieved higher decoding accuracy, whereas the RSA-based approach showed stronger alignment with the target representational structure. This divergence likely reflects the different assumptions underlying the two approaches. Classifier-based approaches emphasize discrete category boundaries and therefore tend to select voxels that maximize separability between labeled classes (Haynes, 2015; Pereira et al., 2009). In contrast, RSA-based approaches model the continuous structure of representational relationships across conditions, capturing graded similarities rather than strict categorical divisions (Kriegeskorte et al., 2008; Kriegeskorte & Kievit, 2013; Walther et al., 2016). The observed spatial differences further align with this contrast, indicating that the two selection approaches may prioritize partially distinct components of the distributed emotion-related network. This distinction is particularly relevant for affective processes, which are commonly described along continuous dimensions and typically evolve gradually rather than shifting abruptly between discrete states. This finding implies that experimental designs employing gradual temporal structure analyses (e.g., temporal RSA), which are commonly applied to time-resolved neural data such as electrophysiological (Chen et al., 2016) or magnetencephalography data (Kolasinski et al., 2020), may benefit from the proposed approach. Furthermore, consistent with this perspective, our empirical results showed that voxels selected by the RSA-based approach retained significant discriminative capacity while maintaining representational alignment with the model structure, whereas voxels selected by the classifier-based approach exhibited weaker correspondence with the structured RDMs. Together, these findings suggest that RSA-based voxel selection may better preserve the relational geometry of emotion representations while remaining sufficiently discriminative, making it well suited for multi-condition cognitive fMRI research. More broadly, these findings raise new questions for future systematic searchlight investigations that move beyond geometrical representations by including topological information (Brown & Farivar, 2025; Lin, 2024; Lin & Kriegeskorte, 2024) within a broader semantic space of orthogonal and non-orthogonal tasks.

### Limitations

A key consideration for the proposed framework is its dependence on the predefined target model RDM. Although using an RDM offers greater flexibility than classification labels which encode only discrete condition boundaries, it also introduces an important limitation: voxel selection becomes highly dependent on the quality of the model RDM itself. Defining an appropriate RDM is not always straightforward, particularly in domains such as emotion where multiple dimensions (e.g., valence, arousal) jointly contribute to the representational structure. In this study, we examined arousal-only, valence-only, and two weighted combined models. While incorporating more complex weightings or additional dimensions may better approximate real-world affective representations, such extensions require stronger theoretical grounding and increase model specification complexity. Future work may benefit from exploring data-driven or adaptive RDMs, as well as hybrid approaches that integrate behavioral measures and computational models to improve the robustness and generalizability of model-based voxel selection.

Another limitation relates to the reliability of voxel responses across repetitions. Reliability-based voxel selection has been shown to improve the stability of multivariate analyses by identifying voxels with consistent responses (Tarhan & Konkle, 2020). Although incorporating a test–retest reliability component into the optimization framework would be desirable, this was not feasible in the current dataset due to the limited number of blocks per run (ten blocks). Splitting the data into odd-even partitions would produce five blocks per split, which is insufficient for generating stable reliability estimates. Under such conditions, adding a reliability assessment would likely introduce additional noise rather than improve voxel selection. Future studies with a larger number of repetitions would allow reliability-based criteria to be more effectively integrated, potentially further enhancing the stability and reproducibility of the selected voxel sets. In addition, using randomized or counterbalanced condition orders in such designs may further help minimize potential sequence-related influences.

A further limitation concerns the performance comparison between the RSA-based and classifier-based voxel selection approaches. In the present study, both approaches were evaluated using their respective optimization metrics (classification accuracy and representational similarity), which cannot directly reflect their relative effectiveness in downstream applications. The feasibility of the RSA-based approach has been demonstrated using the same dataset in our previous work, where participants successfully navigated among multiple emotional states with subject-specific, online-selected voxels (Wang et al., 2026). However, this dataset does not allow a direct offline comparison between the two voxel selection approaches in terms of neurofeedback regulation efficacy. Since regulation performance depends substantially on participant’s learning dynamics and real-time control, retrospective comparisons based on hyperparameters not used during the real-time fMRI neurofeedback experiment will not be valid. To test for potential gains under such settings, future studies should implement different voxel selection approaches in a consecutive way, allowing the searchlight optimization outcome to inform the experimental settings and enabling direct assessment of its impact on neurofeedback performance or other downstream tasks in a controlled way.

## CONCLUSION

This study proposes a subject-level searchlight optimization framework for systematic voxel selection in multi-condition fMRI paradigms. By integrating voxel-wise GLM-based activation analysis with searchlight-based representational similarity analysis, the framework identifies voxel sets that are both task-engaged and sensitive to structured differences among related mental states. A Bayesian optimization–based hyperparameter tuning step further enables efficient identification of high-performing parameter configurations and provides evidence for transferability from pilot to larger samples. Validation using an emotion imagery dataset showed that the multivariate refinement improved alignment between empirical and target representational structures compared with activation-based selection alone. In addition, comparison with a classifier-based voxel selection approach revealed complementary characteristics of the two strategies, with the RSA-based method better preserving structured representational relationships while the classifier-based method achieved higher decoding accuracy. Together, these findings demonstrate that the proposed framework provides an effective and computationally efficient approach for model-informed voxel selection in multi-condition cognitive and affective fMRI research, with potential relevance for individualized characterization of brain function in clinical fMRI studies.

## Notes

### Competing Interest Statement

The authors have declared no competing interest.

### Funding

- RWTH Aachen University Exploratory Research Space (ERS) OPEN Seed Fund
- the START Program of the Faculty of Medicine, RWTH Aachen: 019/23
- RWTH Aachen University Junior Principal Investigator fellowship, Excellence Strategy of the Federal Government and the Laender
- China Scholarship Council: 202206060018

